## Supplemental material for "Arpin deficiency increases actomyosin contractility and vascular permeability"

### Supplemental Figure 1

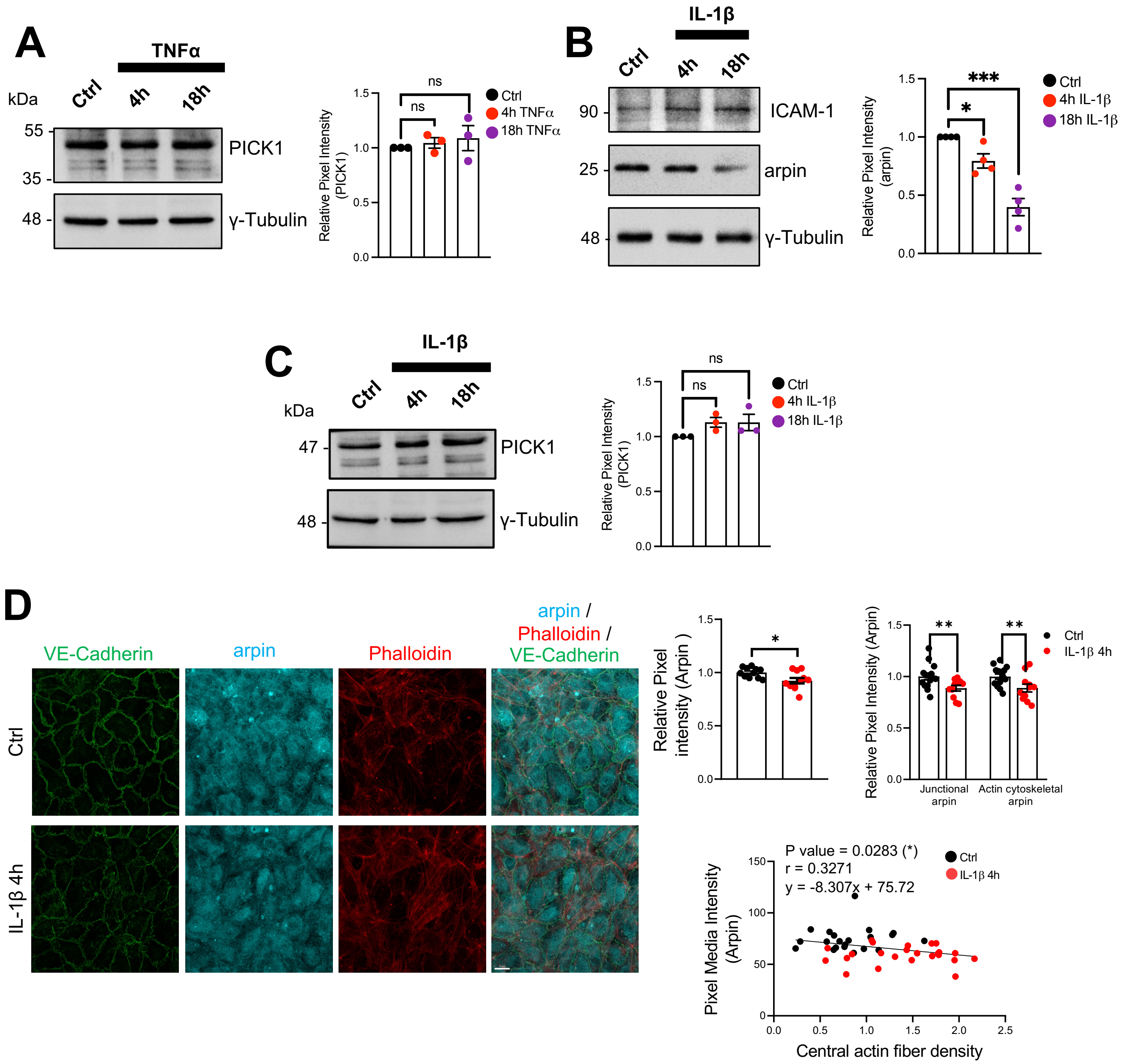

### Supplemental Figure 2

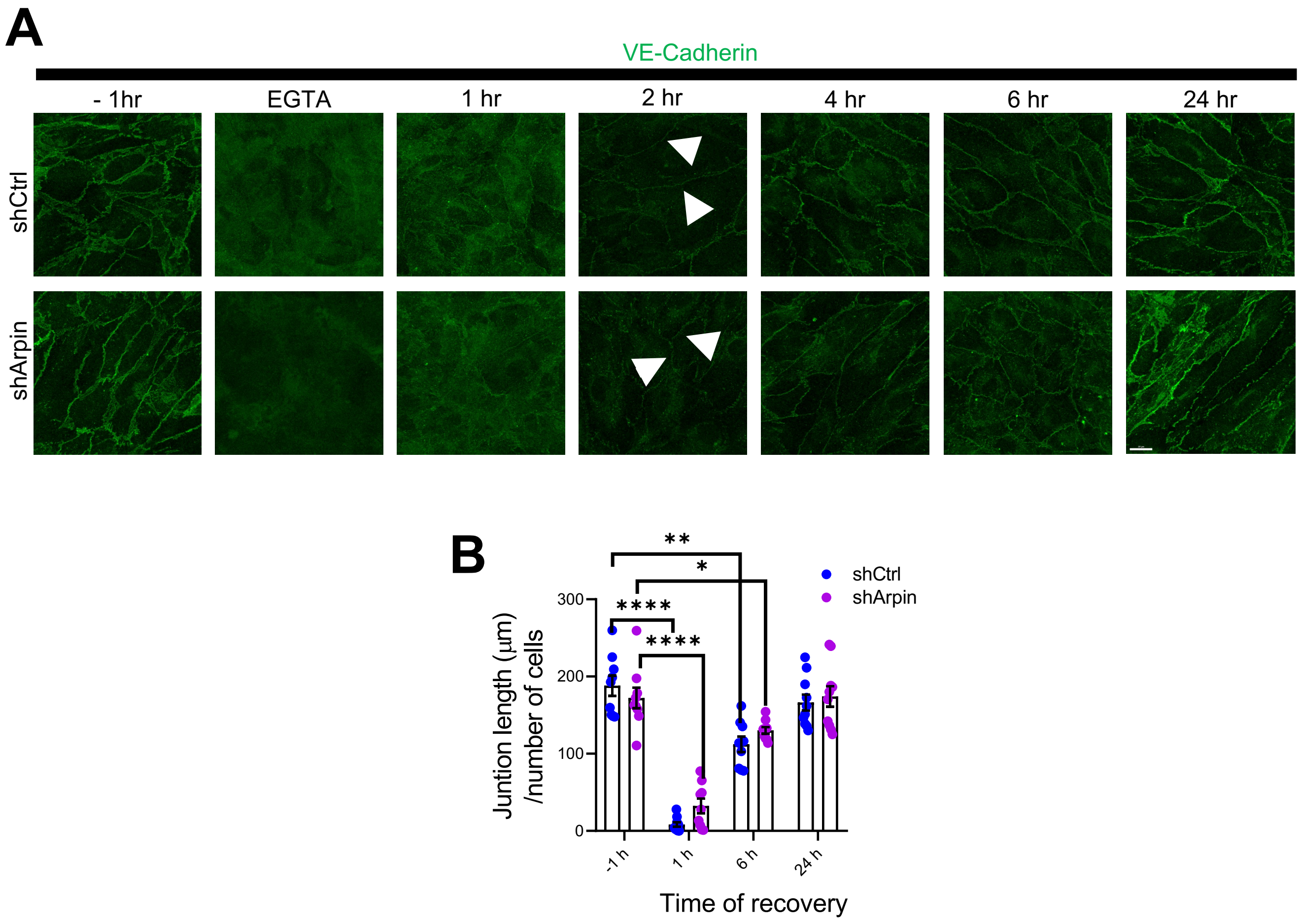

### Supplemental Figure 3

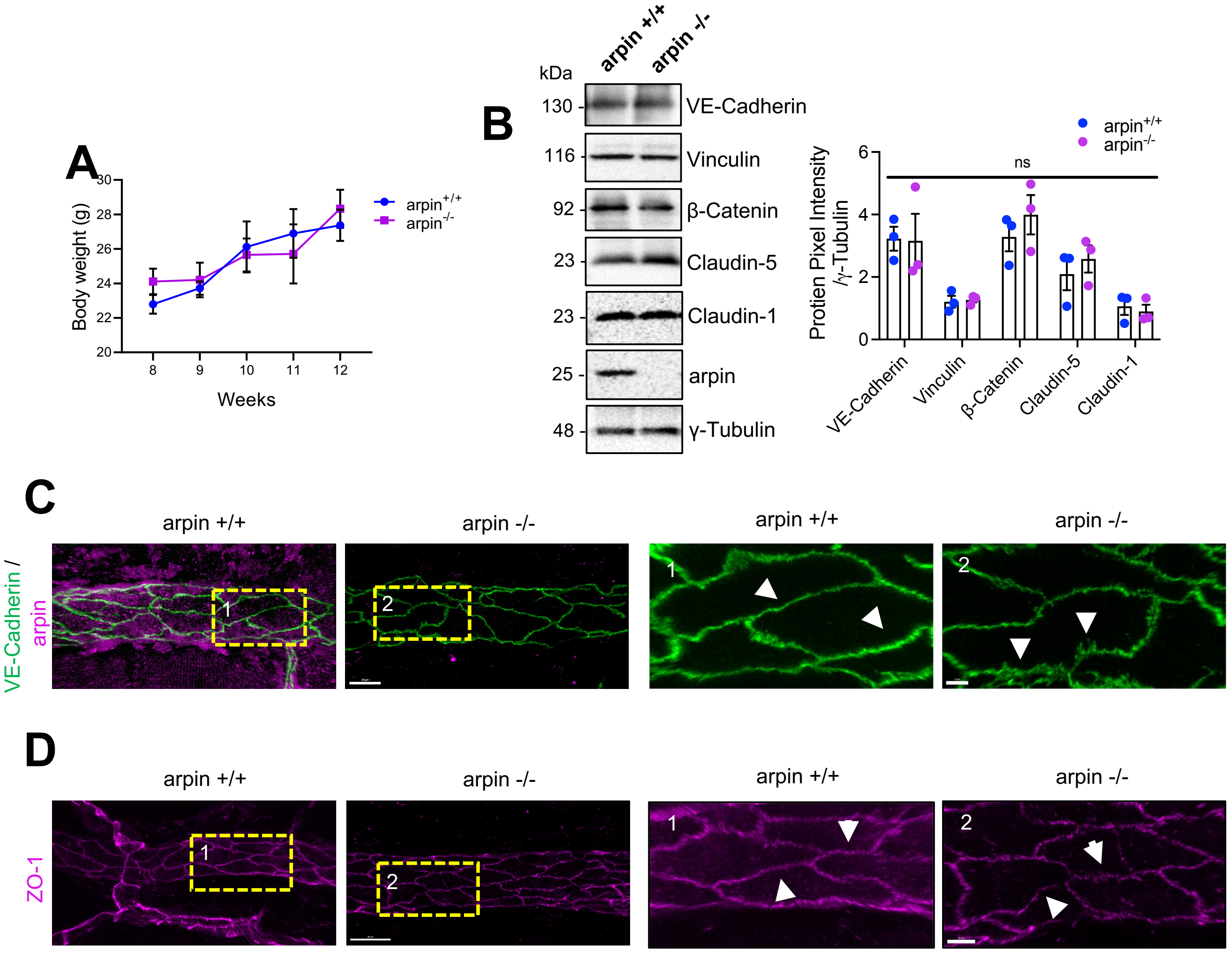

**Supplemental Table S1. Mating statistics of arpin$^{+/-}$ mice**

| | Expected (Theoretical) | Observed (Real) | Probability ($\chi^2$) | Degrees of freedom | P value |
|---|---|---|---|---|---|
| Male | 63 | 66 | 0.286 | 1 | 0.5930 |
| Female | 63 | 60 | | | |
| arpin$^{+/+}$ | 31.5 | 31 | 0.540 | 2 | 0.7635 |
| arpin$^{+/-}$ | 63 | 60 | | | |
| arpin$^{-/-}$ | 31.5 | 35 | | | |

Genotyping of 126 littermates from breedings of heterozygous (arpin$^{+/-}$) mice are shown. $\chi 2$ test probability is shown. Deviations from Mendelian rules are non-significant.

**Supplemental Table S2. Hemograms of arpin$^{+/+}$ and arpin$^{-/-}$ mice.**

| Parameter | arpin$^{+/+}$ (n=8) | | arpin$^{-/-}$ (n=11) | | P value |
|---|---|---|---|---|---|
| | mean | SD | mean | SD | |
| Leukocytes (x10$^{9}$ cells/L) | 3.163 | 1.195 | 2.555 | 1.069 | 0.2599 |
| Neutrophils (x10$^{9}$ cells/L) | 0.787 | 0.790 | 0.527 | 0.349 | 0.3425 |
| Lymphocytes (x10$^{9}$ cells/L) | 2.35 | 1.14 | 1.96 | 0.90 | 0.4085 |
| Monocytes (x10$^{9}$ cells/L) | 0.025 | 0.046 | 0.045 | 0.052 | 0.3897 |
| Erythrocytes (x10$^{12}$ cells/L) | 7.53 | 0.72 | 7.84 | 0.59 | 0.3344 |
| Platelets (x10$^{9}$ cells/L) | 852.0 | 200.3 | 738.1 | 357.3 | 0.4520 |
| Hemoglobin (g/L) | 110.9 | 7.6 | 113.2 | 6.9 | 0.5015 |
| Hematocrit (L/L) | 0.25 | 0.03 | 0.24 | 0.021 | 0.7500 |
